# Gut-resident microorganisms and their genes are associated with cognition and neuroanatomy in children

**DOI:** 10.1101/2020.02.13.944181

**Authors:** Kevin S. Bonham, Guilherme Fahur Bottino, Shelley Hoeft McCann, Jennifer Beauchemin, Elizabeth Weisse, Fatoumata Barry, Rosa Cano Lorente, The RESONANCE Consortium, Curtis Huttenhower, Muriel Bruchhage, Viren D’Sa, Sean Deoni, Vanja Klepac-Ceraj

## Abstract

The gastrointestinal tract, its resident microorganisms, and the central nervous system are connected by biochemical signaling, also known as the “microbiome-gut-brain-axis.” Both the human brain and the gut microbiome have critical developmental windows in the first years of life, raising the possibility that their development is co-occurring and likely co-dependent. Emerging evidence implicates gut microorganisms and microbiota composition in cognitive outcomes and neurodevelopmental disorders (e.g., autism and anxiety), but the influence of gut microbial metabolism on typical neurodevelopment has not been explored in detail. We investigated the relationship of the microbiome with the neuroanatomy and cognitive function of 361 healthy children, demonstrating that differences in gut microbial taxa and gene functions are associated with overall cognitive function and with differences in the size of multiple brain regions. Using a combination of multivariate linear and machine learning (ML) models, we showed that many species, including *Gordonibacter pamelae* and *Blautia wexlerae*, were significantly associated with higher cognitive function, while some species such as *Ruminococcus gnavus* were more commonly found in children with low cognitive scores after controlling for sociodemographic factors. Microbial genes for enzymes involved in the metabolism of neuroactive compounds, particularly short-chain fatty acids such as acetate and propionate, were also associated with cognitive function. In addition, ML models were able to use microbial taxa to predict the volume of brain regions, and many taxa that were identified as important in predicting cognitive function also dominated the feature importance metric for individual brain regions. For example, *B. wexlerae* was the most important species in models predicting the size of the parahippocampal region in both the left and right hemispheres, while several species from the phylum *Bacteroidetes*, including GABA-producing *B. ovatus*, were important for predicting the size of the left accumbens area, but not the right. These findings provide potential biomarkers of neurocognition and brain development and may lead to the future development of targets for early detection and early intervention.

## Introduction

The gut and the brain are intimately linked. Signals from the brain reach the gut through the autonomic nervous system and the endocrine system, and the gut can communicate with the brain through the vagus nerve and through endocrine and immune (cytokine) signaling molecules (*1–4*). In addition, the products of microbial metabolism generated in the gut can influence the brain, both indirectly by stimulating the enteric nervous and immune systems, as well as directly through molecules that enter circulation and cross the blood-brain barrier. Causal links between the gut microbiome and neural development, particularly atypical development, are increasingly being identified (*5*). Both human epidemiology and animal models point to the effects of gut microbes on the development of autism spectrum disorder (*6*, *7*), and specific microbial taxa have been associated with depression (*8*, *9*) and Alzheimer’s disease (*10*, *11*). However, information about this “microbiome-gut-brain axis” in normal neurocognitive development remains lacking, particularly early in life.

The first years of life are critical developmental windows for both the microbiome and the brain (*12*). Fetal development is believed to occur in a sterile environment, but newborns are rapidly seeded at birth through contact with the birth canal (if birthed vaginally), caregivers, food (breast milk or formula), and other environmental sources (*13*, *14*). The early microbiome is characterized by low microbial diversity, rapid succession and evolution, and domination by Actinobacteria, particularly the genus *Bifidobacterium*, Bacteroidetes, especially *Bacteroides*, and Proteobacteria (*15*). Many of these bacteria have specialized metabolic capabilities for digesting human breast milk, such as *Bifidobacterium longum* subsp. *infantis* and *Bacteroides fragilis* (*16*). Upon the introduction of solid foods, the gut microbiome undergoes another categorical transformation; its diversity increases, and most taxa of the infant microbiome are replaced by taxa more reminiscent of adult microbiomes (*13*). Prior studies have typically focused on either infant microbiomes or adult microbiomes, since performing statistical analyses across this transition poses particular challenges. Nevertheless, since this transition coincides with critical neural developmental windows and associated neurodevelopmental processes including myelination, neurogenesis, and synaptic pruning, investigation across this solid-food boundary is important (*17*).

A child’s brain undergoes remarkable anatomical, microstructural, organizational, and functional changes in the first years of life. By age 5, a child’s brain has reached >85% of its adult size, has achieved near-adult levels of myelination, and the pattern of axonal connections has been established (*18*). Much of this development occurs in discrete windows called sensitive periods (SPs) (*19*) during which neural plasticity is particularly high. Emerging evidence suggests that the timing and duration of SPs may be driven in part by cues from the developing gut microbiome (*20*, *21*). As such, understanding the normal spectrum of healthy microbiome development and how it relates to normal neurocognitive development may provide opportunities for identifying atypical development earlier and offer opportunities for intervention.

To begin to address this need, we investigated relationship between the gut microbiome and neurocognitive development in a large cohort of healthy and neurotypically-developing children from infancy through 10 years of age. Gut microbial communities were assessed using shotgun metagenomic sequencing, enabling profiling at both the taxonomic and gene-functional level. Cognitive skills and abilities were measured using age-appropriate psychometric assessments of cognitive function, i.e., the Mullen Scales of Early Learning (MSEL) (*22*), Weschler Preschool and Primary Scale of Intelligence, 4th Edition (WPPSI-4) (*23*) and Weschler Intelligence Scale for Children, 5th Edition (WISC-V) (*24*). Finally, we also assessed emerging brain structure using magnetic resonance imaging (MRI). Through a combination of classical statistical analysis and machine learning, we find that the development of the gut microbiome, children’s cognitive abilities, and brain structure are intimately linked, with both microbial taxa and gene functions able to predict cognitive performance and brain structure.

## Results

### Cohort overview and summary data

We investigated the co-development of the brain and microbiome in 361 healthy and neurotypically-developing children (163 female) between 82 days and 10 years of age (Table 1, Figure 1A, Figure S1) using a variety of orthogonal microbial and neurocognitive measures. These included shotgun metagenomic sequencing, age-appropriate cognitive and behavioral assessments using full-scale composite scores from the MSEL, WPPSI-4 and WISC-V, and neuroimaging measures of cortical and sub-cortical morphology. Different assessment instruments were used depending on the child’s age due to the unique age-range and psychometric properties measured by each (i.e., the MSEL for children under 4 years of age, The WPPSI-3 for 3-6 year-olds, and the WISC-V for children older than 6), but were normalized to a common scale (Figure 1B), As expected, the greatest differences in microbial taxa were observed across child age (Figure 1C-D, PCoA axis 1), with older children primarily stratified into Bacteroidetes-dominant, Firmicutes-dominant, or high-abundance *Prevotella copri* (Figure S2). Overall subject-to-subject variation in gut microbial genes and brain volume profiles was similarly driven largely by subject age (Figure 1E-F).

**Table 1:**
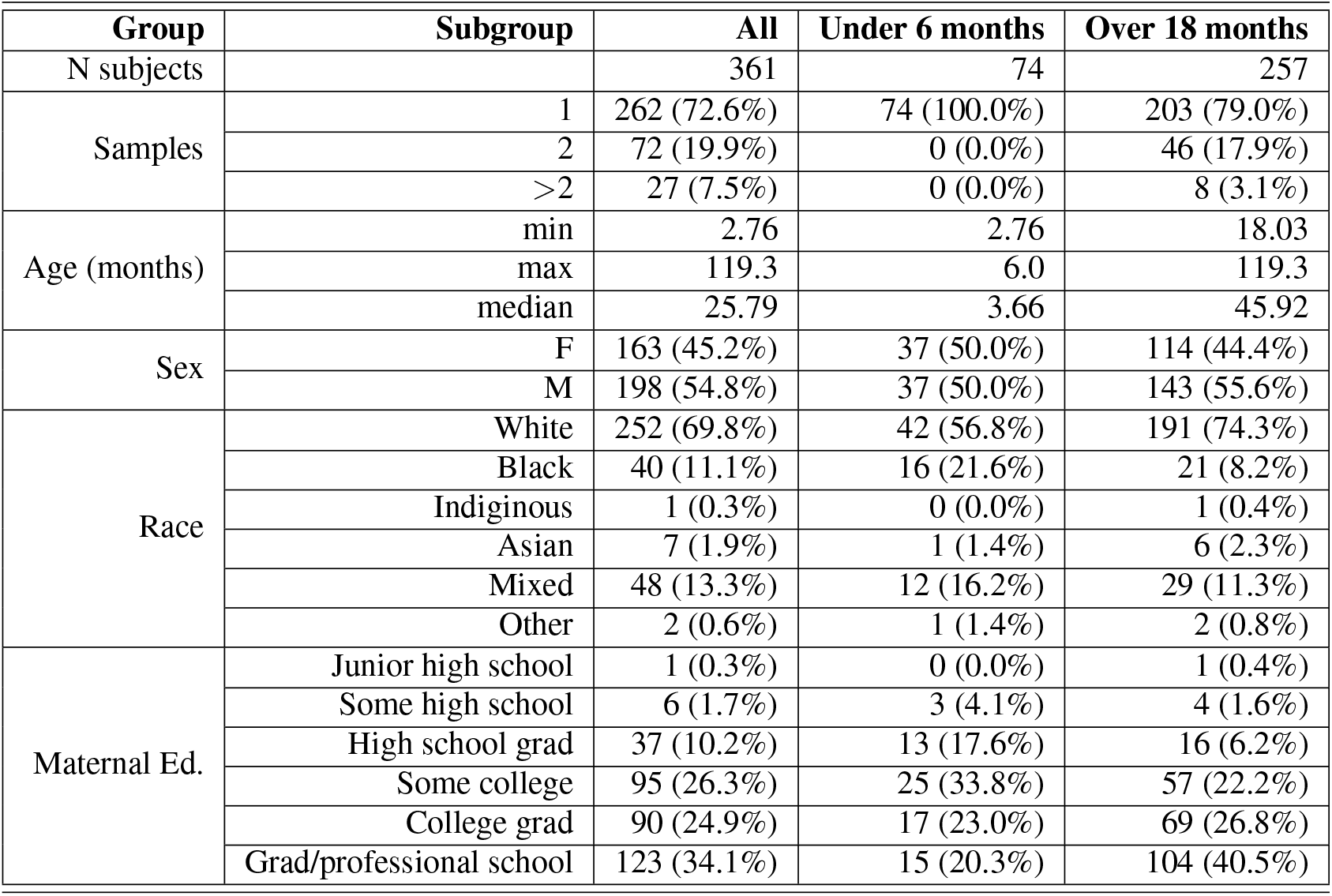
Subject demographics of the ECHO RESONANCE cohort.

**Figure 1:**
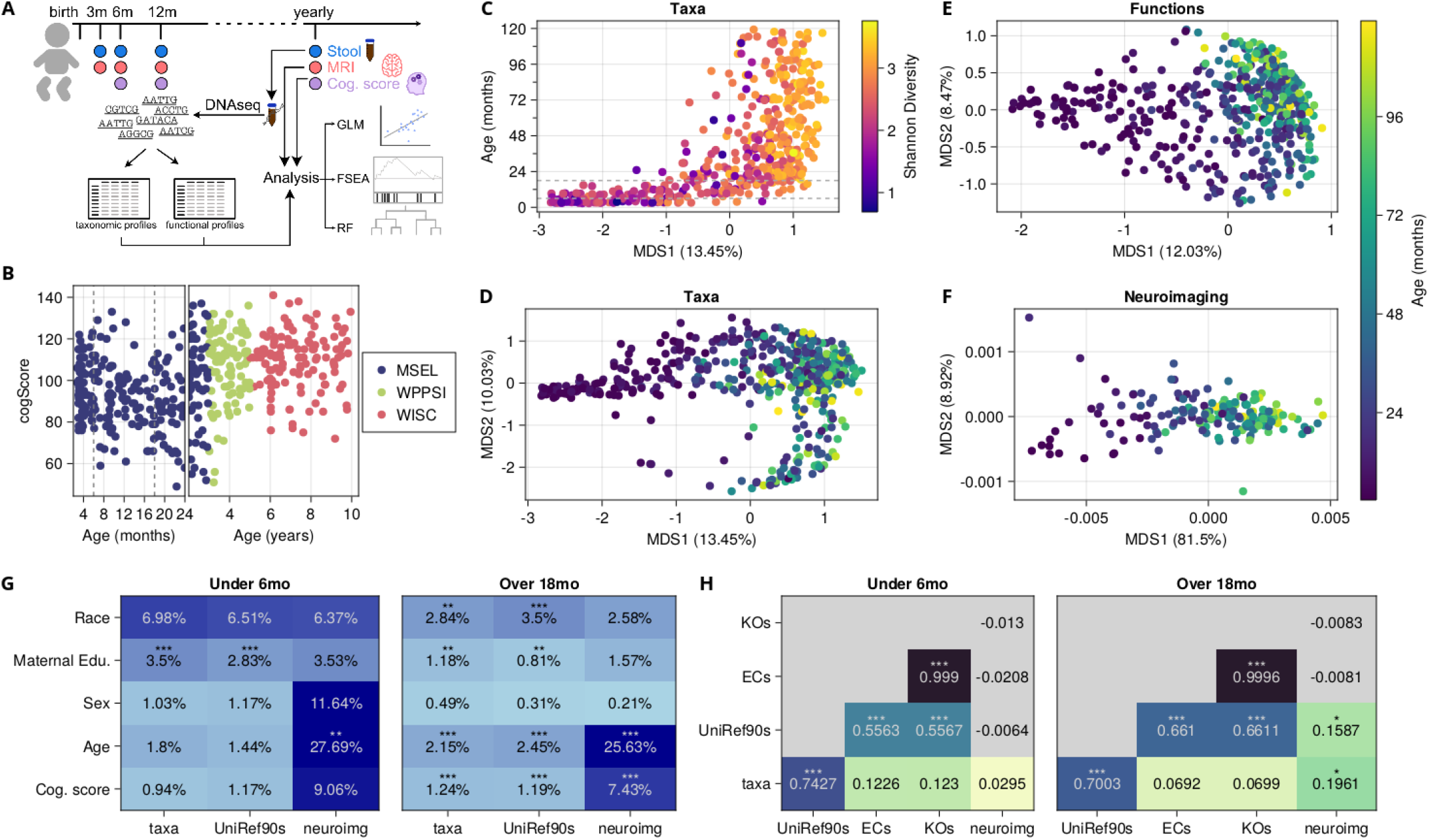
Study design and overview. (A) Stool samples, cognitive assessments and neuroimaging were collected from participants at different ages throughout the first years of life. (B) Cognitive function scores are assessed by different instruments, but may be normalized using full-scale composite scores. **(C-D)** Principal coordinates analysis using Bray-Curtis dissimilarity on taxonomic profiles demonstrates high beta diversity, with much of the first axis of variation explained by increasing age and alpha diversity. Differences in gene function profiles (E) and neuroimaging (PCA based on the Euclidean distance of brain region volumes) **(F)** are likewise dominated by changes as children age. **(G)** Permutation analysis of variance (PERMANOVA) of taxonomic profiles, functional profiles (annotated as UniRef90s) and neuroimaging, against the metadata of interest. Variations in taxonomic and functional profiles explain a modest but significant percent of the variation in cognitive development in children over 18 months of age. (H) Mantel testing of different microbial feature matrices, shows overlapping but distinct patterns of variation. Dotted lines in (B) and (C) show 6 and 18 months, which are used as cut-offs in some following models.

As several prior studies have demonstrated associations between specific gut taxa and atypical neurocognition and neurological disorders (*7*, *8*, *25–27*), we sought to determine if specific taxa or gene functions were associated with normal cognitive development in children. To test if variations in gut microbial taxa, their genes, or their metabolism were associated with neurocognitive development, we used permutation analysis of variance (PERMANOVA) that included the *β* diversity of microbial taxa and gene functions, neuroimaging-derived volume measures of cortical and subcortical structures, and general cognitive ability. Given the large ecological shift in the microbiome that occurs upon the introduction of solid food, and the relatively wide range of ages when over which infant transition from milk to solid foods, we separately considered ages that are generally pre-transition (prior to 6 months of age old) and those that are generally post-transition (over 18 months of age). However, even with this categorization, we find increasing variation in gut microbiomes in children over 18 months old (Figure 1G). We also found that overall variation in microbial species in children over 18 months old was significantly associated with variation in cognitive function score (Figure 1G, R^2^ = 0.0124, q < 0.001), as was variation in microbial gene functions (R^2^ = 0.0119, q < 0.001). Variation in microbial taxa and genes was not significantly associated with cognitive function in children under 6 months, though this may be due to the low taxonomic diversity and broad lack of overlap between taxa in infants (Figure 1G). As expected, age was significantly associated with microbial beta diversity (taxa R^2^ = 0.0215, and gene functions annotated with UniRef90s clusters of 90% similarity (*28*) - R^2^ = 0.0245, q < 0.001), and very strongly associated with variation in neuroimaging profiles (R^2^ = 0.256, q < 0.001).

Consistent with prior studies, different microbial measurement types captured overlapping variation, with species profiles and gene function profiles, both generated from metagenomic sequences, being tightly coupled (Figure 1H, p < 0.001). Other functional groupings (Enzyme commission level4 - ECs (*29*), and KEGG orthologues - KOs (*30*)) overlapped only slightly with taxonomic profiles in both age cohorts, despite being derived from UniRef90 labels. In children over 18 months, some variation (15.9%, p < 0.01) in neuroimaging overlapped with microbial measures, though this may be due to the residual variation due to age in both measures.

### Microbial species and neuroactive genes are associated with cognitive performances

To assess whether individual microbial species were associated with cognitive function, we fit multivariable linear regression models (LMs) (*31*) for the relative abundance of each species that had at least 15% prevalence in a given age group (Figure 2A, N_species_ = 92 for 0–120 months, N_species_ = 46 for 0–6 months, N_species_ = 97 for 18–120 months). Only *Blautia wexlerae* was significantly associated (q value = 0.14, *β* = 0.0015) with cognitive function in children under 6 months old after adjusting for age and maternal education (Figure 2B). By contrast, in children over 18 months of age, several mirobial species were significantly enriched (q-value < 0.20) in children with higher cognitive function scores, including *Gordonibacter pamelaeae, Asaccharobacter celatus* and *Adel-creutzia equolifaciens* (also known as *Adlercreutzia equolifaciens* subsp. *celatus* and *Adlercreutzia equolifaciens* subsp. *equolifaciens* respectively (*32*)), and SCFA-producing probiotic species such as *Eubacterium eligens* and *Faecalibacterium prausnitzii* (*33*) (Figure 2B). *Ruminococcus gnavus* was the only microbial species that was significantly negatively associated with cognitive function score (q value = 0.18).

**Figure 2:**
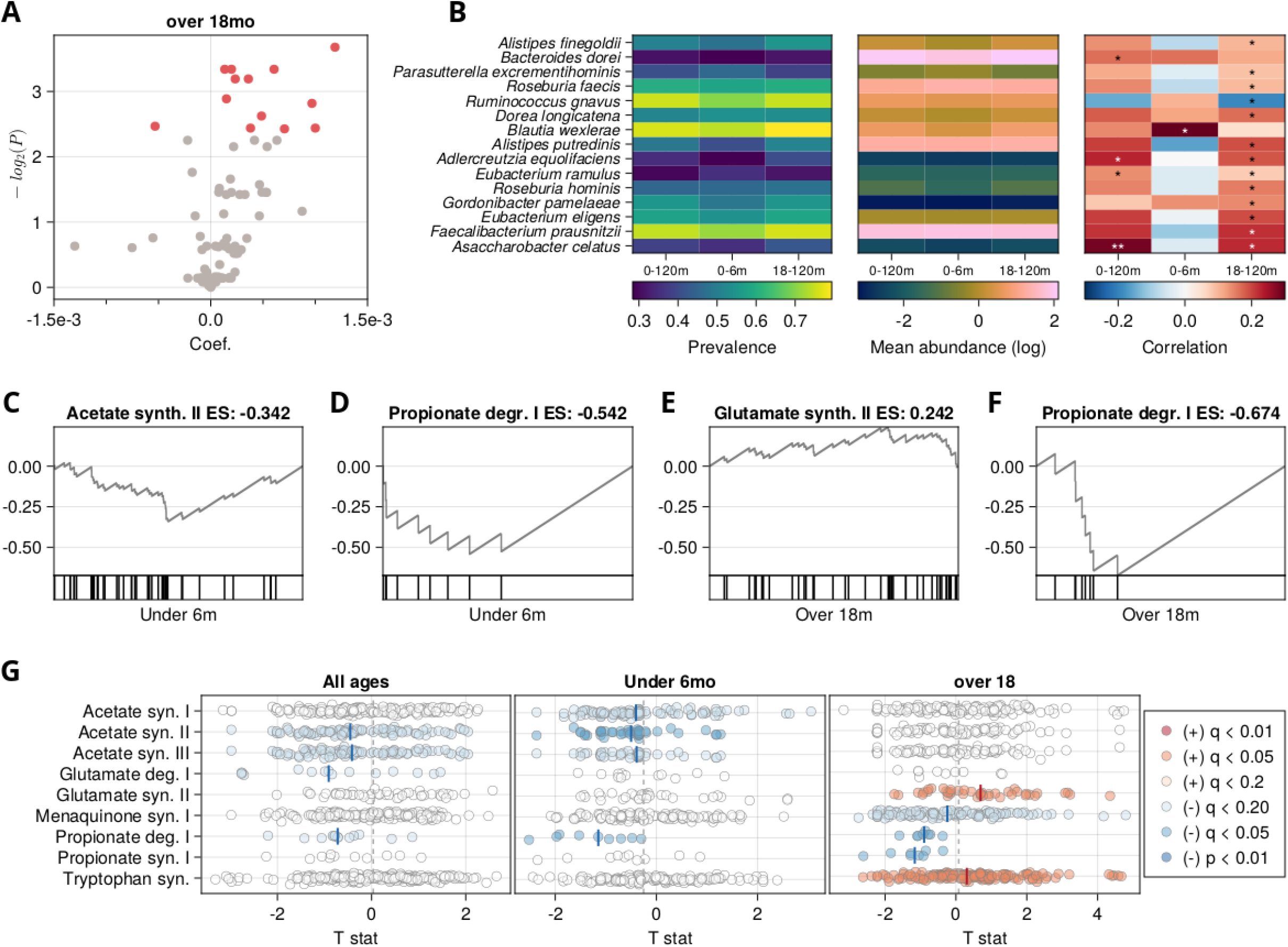
Taxa and gene functional groups are associated with cognitive function. (A) Volcano plot of multivariable linear model results showing the relationship between individual taxa and cognitive function score in children over 18 months of age, controlling for age, maternal education, and sequencing depth. Taxa that were significant after FDR correction (q < 0.2) are colored red. (B) For taxa that were significantly associated with cognitive function, heatmaps of prevalence, mean relative abundance, and correlation with cognitive function in different age groups. **(C-F)** Enrichment plots for selected neuroactive gene sets used in feature set enrichment analysis (FSEA). (G) summary of FSEA results across all samples (left) as well as in the under 6 months (middle) and over 18 months (right) subsets, colored based on the significance of the association. Markers indicate the individual correlation of genes within a gene set, and vertical bars represent the median correlation of that gene set.

Given that different microbial species might occupy the same metabolic niche in different individuals, we hypothesized that microbial genes grouped by functional activity would be associated with cognition. To test this, we performed feature set enrichment analysis (FSEA) on groups of genes with neuroactive potential (*9*) and concurrent cognitive ability score (Table 2, Figure 2C-G) and found that several metabolic pathways were either significantly enriched or depleted in children with higher cognitive ability scores. This was true both when considering all ages (0-120 months) together, though the enrichment of most pathways was more pronounced either in children under 6 months or those over 18 months. For example, genes for degrading the 3-carbon SCFA, propionate, were significantly depleted in children with higher cognitive ability scores across all age groups tested (Table 2; propionate degradation I, under 6 months, enrichment score (E.S.) = −0.542, corrected p-value (q) = 0.020; over 18 months, E.S. = −0.674, q = 0.041). Interestingly, genes for propionate synthesis were also significantly depleted in higher scoring children over 18 months (E.S. −0.676, q = 0.023), as were genes for synthesizing the 2 carbon SCFA acetate in children under 6 months old (acetate synthesis I, E.S. = −0.194, q = 0.153; acetate synthesis II, E.S. = −0.342, q = 0.020; acetate synthesis III, E.S. = −0.31, q = 0.052). In children over 18 months old, synthesis of menaquinone (vitamin K) was also negatively associated with cognitive function score (menaquinone synthesis I, E.S. = −0.170, q = 0.0183), while genes for the synthesis of the amino acids glutamate and tryptophan were significantly enriched in children with higher cognitive function scores (glutamate synthesis I, E.S. = 0.242, q = 0.047; tryptophan synthesis, E.S. = 0.119, q = 0.041). Taken together, these results suggest that microbial metabolic activity, particularly the metabolism (synthesis and degradation) of neuroactive compounds may have effects on cognitive development.

**Table 2:** Feature set enrichment analysis on neuroactive microbial gene sets. Red and blue text indicates significant (q value <0.2) genesets with a positive or negative pseudomedian, respectively.

| Age group | 0 to 120 months |  | 0 to 6 months |  | 18 to 120 months |  |
| --- | --- | --- | --- | --- | --- | --- |
| geneset | E.S. | q value | E.S. | q value | E.S. | q value |
| Acetate syn. I | -0.122 | 0.555 | -0.194 | **0.153** | 0.127 | 0.602 |
| Acetate syn. II | -0.22 | **0.066** | -0.342 | **0.02** | -0.095 | 0.92 |
| Acetate syn. III | -0.21 | **0.086** | -0.31 | **0.052** | -0.086 | 0.92 |
| Glutamate deg. I | -0.404 | **0.18** | -0.284 | 0.816 | -0.249 | 0.602 |
| Glutamate syn. II | 0.095 | 0.602 | -0.134 | 0.98 | 0.242 | **0.047** |
| Menaquinone syn. I | -0.085 | 0.563 | -0.083 | 0.816 | -0.17 | **0.182** |
| Propionate deg. I | -0.461 | **0.13** | -0.542 | **0.02** | -0.674 | **0.041** |
| Propionate syn. I | -0.334 | 0.555 | -0.32 | 0.571 | -0.676 | **0.023** |
| Tryptophan syn. | 0.057 | 0.974 | -0.072 | 0.816 | 0.119 | **0.041** |

### Gut microbial taxonomic and functional profiles predict cognitive function

FSEA relies on understanding functional relationships between individual genes. However, because the relationships between individual taxa are still largely unknown, we turned to Random Forest (RF) models, a family of unsupervised non-parametric machine learning (ML) method that enables the identification of underlying patterns in large numbers of individual features (here, microbial species). Previous studies have reported successful use of RFs for processing highly-dimensional and sparse data from the domain of genomics (*34–38*), along with other works where it was used to predict cognitive conditions related to Alzheimer’s disease in different scenarios (*39*, *40*). Given that gut microbial profiles, as well as neurocognitive development, may partially reflect socioeconomic and demographic factors, we assessed the performance of RF regressors where maternal education (here used as a proxy of socioeconomic status (SES)), sex, and age were included as possible predictors, either alone or in combination with microbial taxonomic profiles (Table 3).

**Table 3:** Benchmark metrics for the cognitive assessment score prediction models. Confidence intervals (C.I.) are calculated from the distribution of metrics from repeated CV at a confidence level of 95. Root mean square error (RMSE) was used to estimate how well the random forest model was able to predict the outcome. Demo. column indicates whether demographic factors such as maternal education, sex and age were included as predictors.

| Subject Ages (months) | Microbial feature | Demo. | Test set RMSE (± C.I.) | Test set correlation (± C.I.) |
| --- | --- | --- | --- | --- |
| 0 to 6 | - | + | 13.01 ± 0.05 | -0.14 ± 0.01 |
| | taxa | - | 12.66 ± 0.05 | -0.1 ± 0.01 |
| | | + | 12.7 ± 0.05 | -0.12 ± 0.01 |
| | genes | - | 12.56 ± 0.05 | -0.01 ± 0.01 |
| | | + | 12.56 ± 0.05 | -0.02 ± 0.01 |
| 18 to 120 | - | + | 16.27 ± 0.04 | 0.506 ± 0.003 |
| | taxa | - | 17.66 ± 0.04 | 0.363 ± 0.003 |
| | | + | 17.29 ± 0.04 | 0.429 ± 0.003 |
| | genes | - | 18.01 ± 0.04 | 0.303 ± 0.003 |
| | | + | 17.86 ± 0.04 | 0.333 ± 0.004 |

As with LMs, RF models for children under 6 months old performed poorly (mean test-set correlation −0.12, mean RMSE 12.70), but RFs were consistently able to learn the relationship between taxa and cognitive function scores in children over 18 months of age (mean test-set correlation 0.429, mean RMSE 17.29). For both age groups, species that were important in RF overlapped with those that were significant when testing the relationship with LMs (Figure 3A-B, Table S1A-B); except for *Eubacterium ramulus*, all taxa that were significant using LMs belonged to a group responsible for 60% of the total importance in RF models (Table 3, Figure 3A). However, the rank-ordering of LM significance and RF importance had substantive differences. For example, in children under 6 months, only *B. wexlerae* was significantly associated with cognitive function in LMs, but was ranked 20th in importance for RF models.

**Figure 3:**
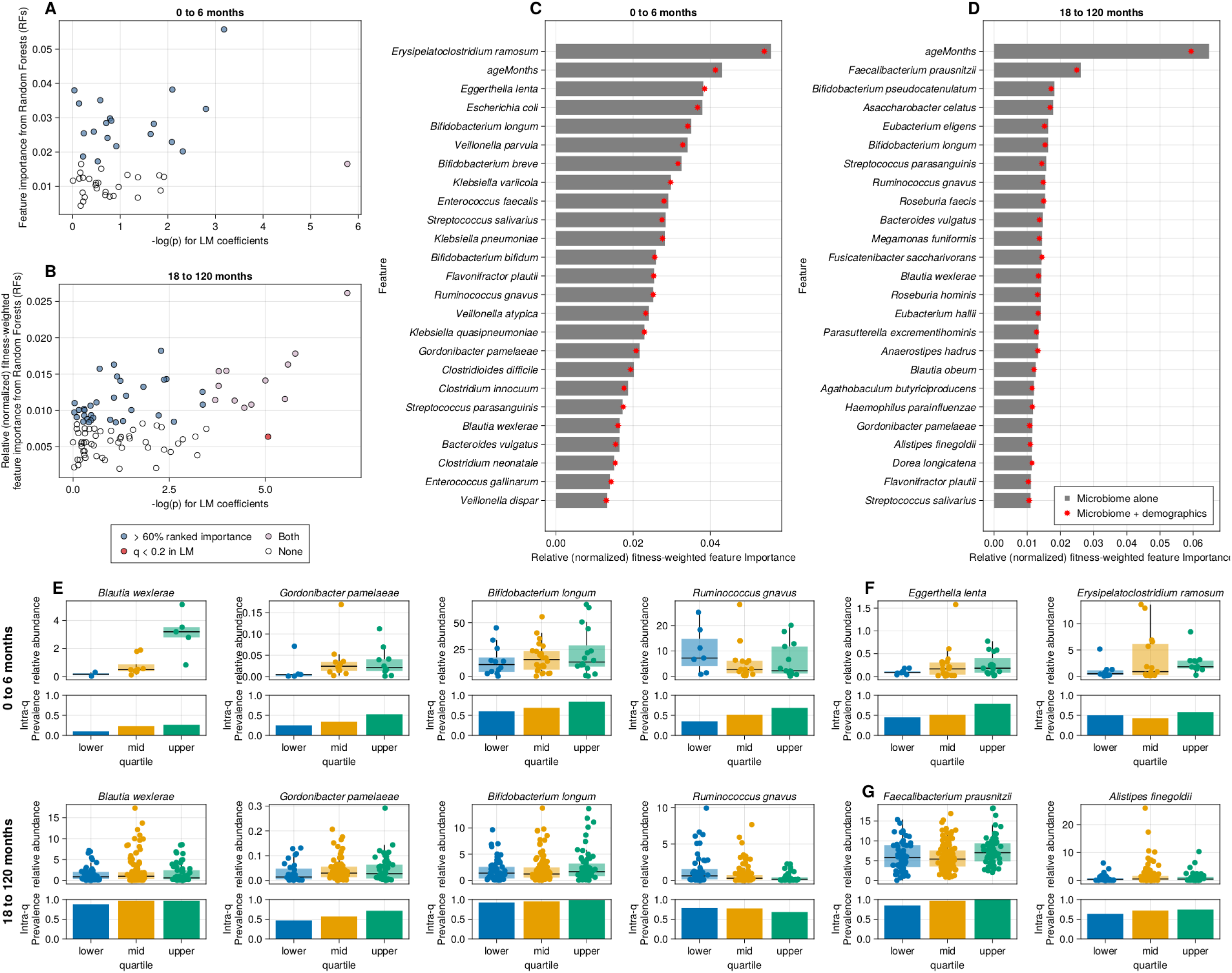
Random forest models predict concurrently measured cognitive function. Comparison of RF predictor importance versus linear models for children between birth and 6 months old **(A),** and for those older than 18 months **(B).** Colors represent whether the species belong to the group of top-important features that account for 60% of the cumulative importance on the RF model, if that species was significant (q < 0.2) in linear models, both, or neither. Ranked feature importance for taxa in RF models for children between birth and 6 months old (C), and for those older than 18 months **(D).** Taxa that are important for RF models both for children under 6 months and for those over 18 months **(E),** only for children under 6 months **(F),** and only for children over 18 months **(G)**

Many taxa highly ranked in importance in RF models in both age groups, including several that were identified in LMs for children over 18 months old, including *R. gnavus* (0–6 months, rank = 13; 18–120 months, rank = 7) and *G. pamelaeae* (0–6 months, rank = 16; 18–120 months, rank = 20), while others such as *Allistipes finegoldii* were age-group specific. A number of taxa that were important in RF models were not statistically significant when using LMs after multiple hypothesis correction. However, these taxa often had small nominal p-values. For example, *Erysipelatoclostridium ramosum* was the most important feature in RF models for children under 6 months old and had an LM p-value of 0.04. Subject age was consistently ranked highly in feature importance, which could indicate that decision branches based on microbial taxa have increased purity when considering the subject’s age or that age itself is a useful predictor.

### Gut microbial taxonomic profiles predict brain structure differences

If there are causal effects of microbial metabolism on cognitive function, they might be reflected in changes in neuroanatomy. We again used RF modeling to associate gut taxonomic profiles with individual brain regions identified in MRI scans (children over 18 months old, N = 121). Some brain regions were more readily predicted by RF models trained on microbial taxa (Table 4, Table S2), in particular those that were highly correlated with age. These included the L/R lingual gyrus (mean RF correlations, Left = 0.421, Right = 0.434; importance for age, Left = 7.5%, Right = 8.3%) and the L/R pericalcarine cortex (mean RF correlations, Left = 0.200, Right = 0.273; importance for age, Left = 3.6%, Right = 6.3%). In many cases, however, age was not an important variable in high-performing models, such as that for the left accumbens area (mean RF correlations = 0.288; relative age importances = 1.2%).

**Table 4:** Summary statistics for the neuroanatomy prediction benchmarks. Mean absolute proportional error (MAPE) and correlation coeficient (R) of test-set preditions are reported accross models for all brain regions. Complete list of values can be found on Table S2.

| Statistic | MAPE | R |
| --- | --- | --- |
| Mean | 0.073 | 0.137 |
| Standard deviation | 0.028 | 0.137 |
| Maximum | 0.206 | 0.605 |
| 75th percentile | 0.082 | 0.195 |
| 50th percentile | 0.067 | 0.134 |
| 25th percentile | 0.057 | 0.062 |
| Minimum | 0.035 | -0.147 |

While models for many regions were highly concordant across the left and right hemispheres in terms of both model performance and microbial feature importance, other regions had substantial bilateral differences. For example, the left accumbens area has one of the highest test-set correlations of our brain region models (R = 0.288), as compared to the right which had a negative mean test-set correlation (R = −0.041), indicating that generalizable models were not identified. Feature importance for models of the left accumbens area were dominated by three species of *Bacteroides, B. vulgatus* (3.3% importance), *B. ovatus* (3.7% importance), and *B. uniformis* (2.9% importance).

While RF models for multiple brain regions had many important microbial taxa, others were dominated by a small number of taxa, many of which were also identified in LMs and RF models of cognitive function (Figures 2 and 3). In general, for all segments, a consistent number of species between 31 and 45 (median = 37, less than 30% of the input features) was responsible for half of the cumulative importance, regardless of model performance (Figure 4A-B, Table S2). We observed two major patterns of importance distribution from species over the brain segments; some species portrayed high contributions to multiple different segments, while others contributed modestly to just one or two brain segments. Notable cases of the first pattern included seven species - *Anaerostipes hadrus*, *B. vulgatus*, *Fusicatenibacter saccharivorans*, *Ruminococcus torques*, *Eubacterium rectale*, *Coprococcus comes* and *B. wexlerae*, which combined, account for approximately 10% of the cumulative relative importances, computed after fitness-weighting on the reported brain segments (Figure 4A-B).

**Figure 4:**
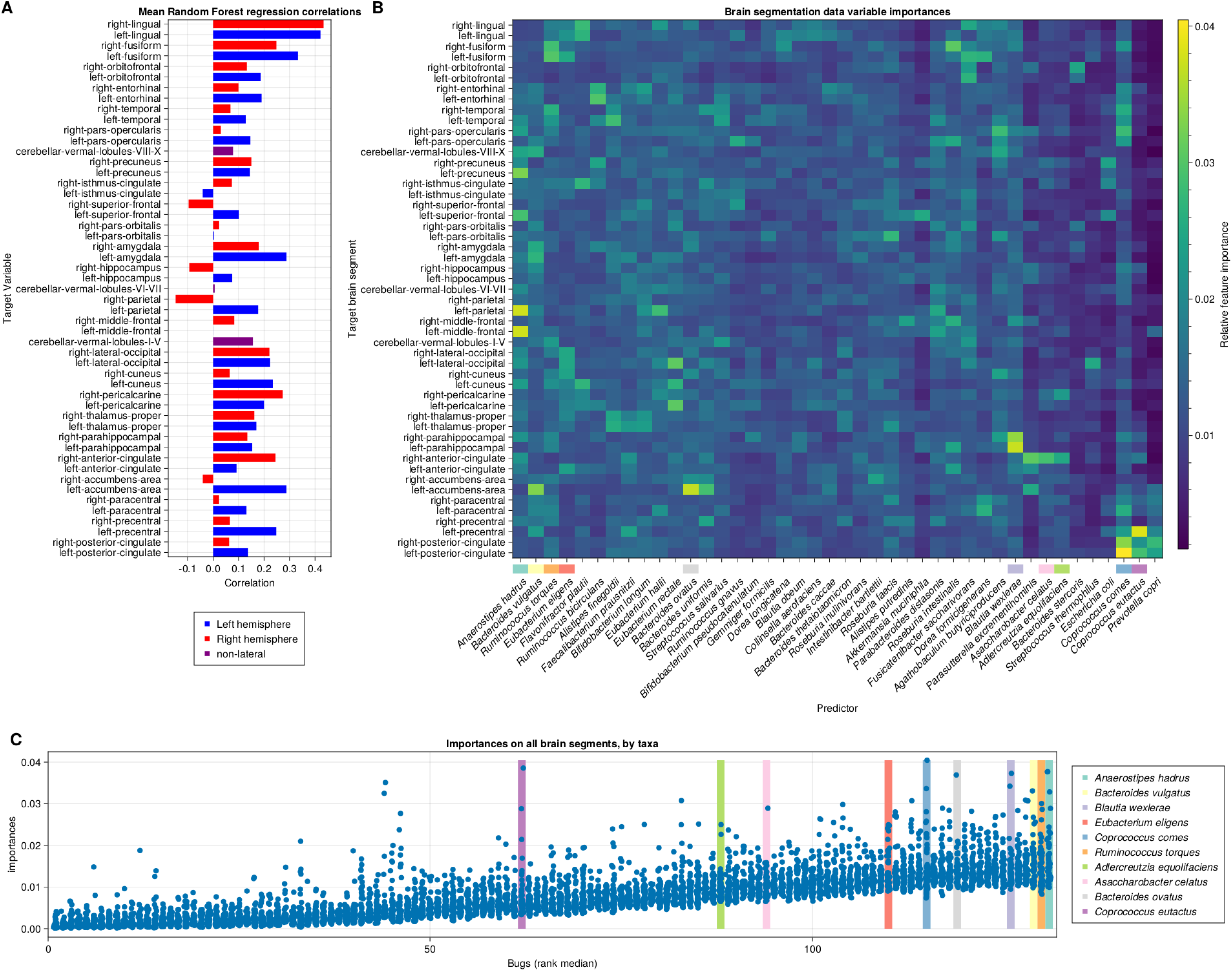
Microbial feature importance in predicting brain volumes in children over 18 months of age. **(A)** Average test-set correlations for prediction of MRI segmentation data from microbiome and demographics on select segments after repeated cross validation. **(B)** Heatmap of average individual relative taxa importances on each brain segment. Importances are reported as proportions relative to the sum of importances for each model - since every model is trained with 132 features, and even distribution of importance would be 0.75% for each feature. Segments and taxa ordered by HCA on a select list of species with high ranks on average importances, and their respective highest-load segments **(C).**

Among these, *A. hadrus* is the variable with highest average relative importance along all reported (mean = 1.7%; min = 1.0%; max = 3.8%, Figure 4B-C). *B. wexlerae*, which was important in RF models of cognitive function, and was the only species to be significantly associated with cogntive function in young children using LMs, was also important in models of the entorhinal, fusiform and lingual areas and had the highest importance in models predicting the size of both the left and right right parahippocampus. Another cluster of model importance contained taxa associated with the basal forebrain, especially the cingulate and the accumbens. This cluster contains *B. ovatus* and *B. uniformis*, but also *Alistipes finegoldii and Streptococcus salivaris*.

Both *A. equolofaciens* and *A. celatus*, two closely-related equol-producing species, are examples of the second contribution pattern, where a taxon only contributes significantly to a very small number of segments. Each of these species were highly important in predicting the relative volume of the right anterior cingulate (respectively, 3.0% and 2.6% importances). Another example of this pattern is *R. gnavus*, which was the only species with significant negative association with cognitive function in linear models. It is heavily associated with the left pars opercularis (2.7% importance). Finally, *C. comes* has an importance distribution that splits almost evenly among the two previously reported clusters. Its loading on the prediction of the left posterior cingulate is the highest for a microbe in RF models (4.0% importance); only age had higher relative importance in any model. At the same time, *C. comes* is one of the most important predictors for neighboring areas of the posterior cingulate such as the pars opercularis (importances, Left = 2.3%, Right = 2.8%), along with upper regions like the left precentral (2.3% importance) and paracentral lobes (importances, Left = 2.6%, Right = 1.9%).

## Discussion

The relationship between the gut microbiome and brain function via the gut-microbiome-brain axis has gained increasing acceptance largely as a result of human epidemiological studies investigating atypical neurocognition (eg anxiety and depression, neurodegeneration, attention deficit / hyperactivity disorder, and autism (*41*, *42*)) and mechanistic studies in animal models (*27*, *43*). The results from these studies point to the possibility that gut microbes and their metabolism may be causally implicated in cognitive development, but this study is the first to our knowledge that directly investigates microbial species and their genes in relation to typical development in young, healthy children. Understanding the gut-brain-microbiome axis in early life is particularly important, since differences or interventions in early life can have outsized and longer-term consequences than those at later ages due to the dynamic and plastic nature of both the gut microbiome and the brain. Further, even in the absence of causal impacts of microbial metabolism, identifying risk factors that could point to other early interventions would also have value.

We identified several species in the family Eggerthelaceae that were associated with cognitive function, including *G.pamelaeae, A. equolofaciens, A. celatus* (formerly regarded a subspecies of *A. equolofaciens* (*44*)), and *Eggerthella lenta*. Many members of this family are known in part due to unique metabolic activities. For example, *A. equolofaciens* produces the nonsteroidal estrogen equol from isoflaven found in soybeans (*45*). This is particularly intriguing, since both *A. equolofaciens* and *A. celatus* were also the most important features in RF models of the right anterior cingulate, which has been linked to social cognition and reward-based decision making (*46–48*), and estrogen has previously been shown to have activity in this brain region (*49*). *G. pamelaeae* also has a unique metabolism that may impact the brain, as it can metabolize the polyphenol ellagic acid (found in pomegranates and some berries) into urolithin, which has been shown in some studies to have a neuroprotective effect (*50*, *51*). While we could not find published evidence of neurological roles for *E. lenta*, it has been extensively studied for its ability to metabolize drug compounds such as the plant-derived heart medication digoxin (*52*). The metabolic versatility of this clade and the large number of species that are associated with cognition make these microbes prime targets for further mechanistic studies.

Other bacteria that were associated with both cognitive ability and brain structure had also have possible connections to previous studies. For example, RF models predicting the size of the left nucleus accumbens were dominated by *Bacteroides* species, especially *B. vulgatus*, *B. uniformis*, and *B. ovatus*. *B. ovatus* has been shown to produce a number of neuroactive compounds including SCFAs and neurotransmitters such as GABA (*53*). Another *Bacteroides* species, *B. fragilis* was able to ameliorate ASD-like symptoms in a mouse model by consuming a microbially-dependent metabolite 4-ethylphenylsulfate (4EPS) (*43*), while *B. ovatus* is a primary producer of a precursor to 4EPS (*27*) and increased anxiety behaviors in mice. The fact that the importance of these species in RF models, and the high performance of RF models in predicting this region were unilateral (affecting the left but not the right accumbens area) was also notable. Most healthy individuals have a rightward asymmetry in the nucleus accumbens, which is involved in reward learning and risk taking behaviours (*54*, *55*). Reduced asymmetry between right and left nucleus accumbens has been associated with substance use disorder in young adults (*56*). The accumbens area is also associated with reward control, and in individuals diagnosed with ADHD, it has been shown to have a divergent neuromorphology (*57*).

As is made apparent by the potentially opposing roles of *B. fragilis* and *B. ovatus*, individual species from the same genus may play different roles. As such, the use of shotgun metagenomic sequencing may be crucial to identifying microbial influences on the brain, since it enables species-level resolution of microbial taxa. A previous study of cognition in 3 year old subjects used 16S rRNA gene amplicon sequencing and showed that genera from the Lachnospiraceae family as well as unclassified Clostridiales (now Eubacteriales) were associated with higher scores on the Ages and Stages Questionnaire (*58*). However, each of these clades encompass dozens of genera with diverse functions, each of which may have different effects. Indeed, several of the taxa that were positively associated with cognitive function in this study, including *B. wexlerae, D. longicatena, R. faecis*, and *A. finegoldii* are Clostridiales, as is *R. gnavus*, which we found was negatively associated with cognitive function (Figure 2A-B). This kind of species-level resolution is typically not possible with amplicon sequencing.

In addition to improved species-level resolution, shotgun metagenomic sequencing also enables gene-functional insight. We showed that genes for the metabolism of SCFAs, both their degradation and synthesis, are associated with cognitive function scores (Figure 2). SCFAs are produced by anaerobic fermentation of dietary fiber and have been linked with immune system regulation as well as directly with brain function (*59*). Microbial metabolism of other potentially neuroactive molecules were also associated with cognitive ability. For example, we were particularly interested in genes for metabolizing glutamate, which is a critical neurotransmitter controlling neuronal excitatory/inhibitory signaling along with gamma-aminobutyric acid (GABA), and their balance in the brain controls neural plasticity and learning, particularly in the developing brain (*60*, *61*). However, while the differential abundance of genes for the metabolism of neuroactive compounds like these is suggestive, it is difficult to reason about the relationship between levels of these genes and the gut concentrations of the molecules their product enzymes act on since the presence or absence of metabolites may select for microbes with the ability to degrade or produce that metabolite respectively. For this reason, future studies coupling shotgun metagenomics with stool metabolomics could improve our understanding of the relationship between microbial metabolism and cognitive development. Further, strain-level analysis linking specific gene content in species of interest could further refine targeted efforts at identifying specific metabolic signatures of microbe-brain interactions.

The use of multiple age-appropriate cognitive assessments that could be normalized to a common scale enabled us to analyze microbial associations across multiple developmental periods, but carries several drawbacks. In particular, the test-retest reliability, as well as systematic differences between test administrators may introduce substantial noise into these observations, particularly in the youngest children. In addition, our study period overlapped with the beginning of the COVID-19 pandemic, and we and others have observed some reduction in measured scores for children that were assessed after the implementation of lockdowns. In our subject set for this study, these effects are more pronounced in some age groups due to the period when active recruitment for the study was occurring (*62*, *63*) (Figure S3).

This analysis allowed us to establish links between microbial taxa and their functional potential with cognition and brain structure. Although we cannot test causality or the chemistry behind the interactions between gut microbial taxa and their genes, the gut, and the brain, this study provides clear and statistically significant associations between the infant and early child gut microbiota and neurocognition. Future studies should focus on characterizing the early-life microbiome and neurocognitive development across different geographic regions and lifestyles such as covering traditionally understudied populations, such as low-resource urban, peri-urban and rural communities to obtain the more comprehensive understanding of the variability within the different gut microbiomes reflects on neurocognition. These studies would also provide us with the wealth of data on different strains from the same species to better understand the effect of genes and their products. Furthermore, culturing and microbial community enrichment studies combined with genetic manipulation and genomic approaches to understand microbial metabolism at the molecular level is key, as the metabolic functions shape and influence the human host and its health. The discovery of the neuroactive metabolites could provide us with biomarkers for early detection or necessary medicinally useful molecules that can be applied in intervention.

## Material and methods

### Study Ethics

All procedures for this study were approved by the local institutional review board at Rhode Island Hospital, and all experiments adhered to the regulation of the review board. Written informed consent was obtained from all parents or legal guardians of enrolled participants.

### Participants

Data used in this study were drawn from an ongoing longitudinal brain and cognitive development, termed RESONANCE, based in Providence, RI, USA, and spanning fetal through adolescent development. The RESONANCE cohort is part of the NIH Environmental influences on Child Health Outcomes (ECHO) study (*64*, *65*), and includes neuroimaging (magnetic resonance imaging, MRI), neurocognitive assessments, bio-specimen analyses, subject genetics, broad environmental exposure data (such as chemical, water quality, nutrition, etc.), and rich demographic, socioeconomic, family and medical history information.

As a study of neurotypical development, mothers and their children were devoid of major risk factors for abnormal or atypical development. These included preterm birth (<37 weeks gestation) low birth weight (<1500g), in utero exposure to alcohol, cigarette or illicit substance exposure; abnormalities on fetal ultrasound; non-singleton or complicated pregnancy, including preeclampsia, high blood pressure, or gestational diabetes; complicated vaginal or cesarian birth; 5 minute APGAR scores <8; NICU admission; neurological disorder in the child (e.g., head injury resulting in loss of consciousness, epilepsy); and psychiatric or learning disorder (including maternal depression) in the infant, parents, or siblings requiring medication in the year prior to pregnancy. In addition to screening at the time of enrollment, on-going screening for worrisome behaviors using validated tools was performed to identify at-risk children and remove them from subsequent analysis. For this study, 361 typically-developing children between the ages of 2.8 months and 10 years old (median age 2 years, 2 months) were selected for analysis based on having provided at least one stool sample in the same week as being evaluated using one of the cogntitive ability assessments (MSEL, WPPSI, WISC-V).

### Cognitive Assessments

General cognitive ability was assessed using age-appropriate performance-based measures. For children under 3 years of age, we used the Early Learning Composite (ELC) from the Mullen Scales of Early Learning (MSEL) (*22*). The MSEL isa standardized and population-normed tool that assesses function in 5 major domains: fine and gross motor, expressive and receptive language, and visual functioning. The ELC is an age-normalized composite derived from these individual domains with a mean of 100 and standard deviation of 15. For older children, full-scale IQ (FSIQ) was assessed using the Wechsler Preschool and Primary Scale of Intelligence 4th Edition (WPPSI-IV, children 3 to 6) (*23*), and the Wechsler Intelligence Scale for Children 5th Edition (WISC-V) for children older than 6 (*24*). Like the ELC, FSIQ is normalized to a mean of 100 and standard deviation of 15.

### Stool Sample Collection and Sequencing

Stool samples (n = 493) were collected by parents in OMR-200 tubes (OMNIgene GUT, DNA Genotek, Ottawa, Ontario, Canada), stored on ice, and brought within 24 hrs to the lab in RI where they were immediately frozen at −80°C. Stool samples were not collected if the subject had taken antibiotics within the last two weeks. DNA extraction was performed at Wellesley College (Wellesley, MA). Nucleic acids were extracted from stool samples using the RNeasy PowerMicrobiome kit, excluding the DNA degradation steps. Briefly, the samples were lysed by bead beating using the Powerlyzer 24 Homogenizer (Qiagen, Germantown, MD) at 2500 rpm for 45 s and then transferred to the QIAcube (Qiagen, Germantown, MD) to complete the extraction protocol. Extracted DNA was sequenced at the Integrated Microbiome Resource (IMR, Dalhousie University, NS, Canada).

Shotgun metagenomic sequencing was performed on all samples. A pooled library (max 96 samples per run) was prepared using the Illumina Nextera Flex Kit for MiSeq and NextSeq from 1 ng of each sample. Samples were then pooled onto a plate and sequenced on the Illumina NextSeq 550 platform using 150+150 bp paired-end “high output”;’ chemistry, generating 400 million raw reads and 120 Gb of sequence per plate.

### Computational analysis of metagenomes

Shotgun metagenomic sequences were analyzed using the bioBakery suite of computational tools (*66*). First, KneadData (v0.7.7) was used to perform quality control of raw sequence reads, such as read trimming and removal of reads matching a human genome reference. Next, MetaPhlAn (v3.0.7, using database mpa_v30_CI I()C()PhlAn_201901) was used to generate taxonomic profiles by aligning reads to a reference database of marker genes. Finally, HUMAnN (v3.0.0a4) was used to functionally profile the metagenomes.

### Machine learning for cognitive development

Prediction of cognitive scores was carried out as a set of regression experiments targeting real-valued continuous assessment scores. Different experiment sets were designed to probe how different representations of the gut microbiome (taxonomic profiles, functional profiles encoded as ECs) would behave, with and without the addition of demographics (sex and maternal education as a proxy of socioeconomic status) on participants from different age groups. Age (in months) was provided as a covariate for all models (Table S3).

Random Forests (RFs) (*67*) were selected as the prediction engine and processed using the DecisionTree.jl (*68*) implementation, inside the MLJ.jl (*69*). Independent RFs were trained for each experiment, using a set of default regression hyperparameters from Breiman and Cutler (*67*), on a repeated cross-validation approach with different RNG seeds. One hundred repetitions of 3-fold CV with 10 different intra-fold RNG states each were employed, for a total of 3000 experiments per input set.

After the training procedures, the root-mean-square error (RMSE) for cognitive assessment scores and mean absolute proportional error (MAPE) for the brain segmentation data, along with Pearson’s correlation coefficient (R) were benchmarked on the validation and train sets. MAPE was chosen as the metric for brain segments due to magnitude differences between median volumes of each segment, which would hinder interpretation of raw error values without additional reference.

To derive biological insight from the models, the covariate variable importances for all the input features, measured by mean decrease in impurity (MDI, or GINI importance), was also analyzed. Leveraging the distribution of results from the extensive repeated cross validation experiments, rather than electing a representative model or picking the highest validation-set correlation, a measure of model fitness (**Equation 1**) was designed to weight the importances from each trained forest, and all ‘‘importance” values reported refer to the average fitness-weighted importance accross all models. The objective was to give more weight to those with higher benchmarks on the validation sets (or higher generalizability), while penalizing information from highly overfit models, drawing inspiration from the approach used on another work employing repeated CV on Random Forests with high-dimensional, low sample size datasets (*70*). The resulting fitness-weighted importances were used to generate the values in Figures 3–4.

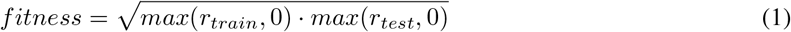

### MRI / segmentation

T1-weighted anatomical MRI data were acquired using a 3-Dimensional MP-RAGE sequence on 3T Siemens Trio scanner with the following parameters: TE=5.6msec, TR=1400msec, TI = 950msec, FA=15 degrees, and 1.1×1.1×1.1mm resolution. Image matrix and field-of-view were adjusted by infant size to achieve the desired image resolution. For children under 4 years of age, MRI was performed during natural non-sedated sleep using previously described methods (*71*). Older children were scanned awake whilst watching a movie or favorite TV show. To help minimize motion, pediatric (MedVac) immobilizers were used along with foam cushions and padding. Image quality was monitored during and following scanning. If motion artifacts were observed (e.g., ghosting, blurring, or other signal variations), scans were repeated or removed from analysis.

Following data acquisition, brain region volumes corresponding to total white and gray matter, cortical gray matter, brainstem, and left and right hemisphere lateral ventricles, thalamus, caudate, putamen, pallidum, hippocampus, amygdala, and nucleus accumbens were calculated using an atlas matching approach (*72*). Individual images were first nonlinearly aligned to MNI-space using a multi-step registration approach (ANTs 2.2 toolbox (*73*)). The MNI-aligned Harvard-Oxford brain atlas was then aligned and superimposed onto the individual child by applying the inverse transformation, and the brain regions of interest delinated (segmented) and their volume calculated by summing the number of image voxels within each (*74*). Brain region volumes used in RF models were normalized to total brain volume (the sum of white and gray matter).

## Data and code availability

All data needed to evaluate the conclusions in the paper are present in the paper and/or the Supplementary Materials. Taxonomic and functional microbial profiles, as well as subject demographics necessary for statistical analyses and machine learning are available on the Open Science Framework (*75*). Data processing, generation of summary statistics, and generation of plots was performed using the julia programming language and 3rd party packages (*69*, *76–81*). Information for replicating the package environment as well as code for data analysis and figure generation, as well as scripts for automated download of input files are available on GitHub (*82*).

## Acknowledgements

This research was funded by NIH UG3 OD023313 and Wellcome LEAP 1kD. The authors declare that they have no competing interests.

KSB, VKC, CH, and SD were responsible for conceptualization, methodology and project administration. VKC, SD, and VD were responsible for funding aqusition. KSB, SHM, MB, FB, RCL, and JB contributed to data curation. Formal analyses were conducted by KSB, GFB, and MB. Investigation was performed by KSB, GFB, SHM, EW, FB, RCL, and MB. Resources were provided by SHM, EW, FB, RCL, JB, CH, and MB. Software and data visualizations were produced by KSB and GFB. KSB, MB, SD, VD, and VKC were responsible for supervision. Validation was performed by KSB, SHM, JB, and MB. KSB and GFB wrote the initial draft, and review and editing were provided by SD, CH, and VKC.

## Notes

https://osf.io/ybs32/

https://zenodo.org/record/7647510

### Competing Interest Statement

The authors have declared no competing interest.

### Summary of Updates

Minor edits, typos, figure style

