## Supplemental Tables and Figures for "Gut-resident microorganisms and their genes are associated with cognition and neuroanatomy in children"

March 31, 2023

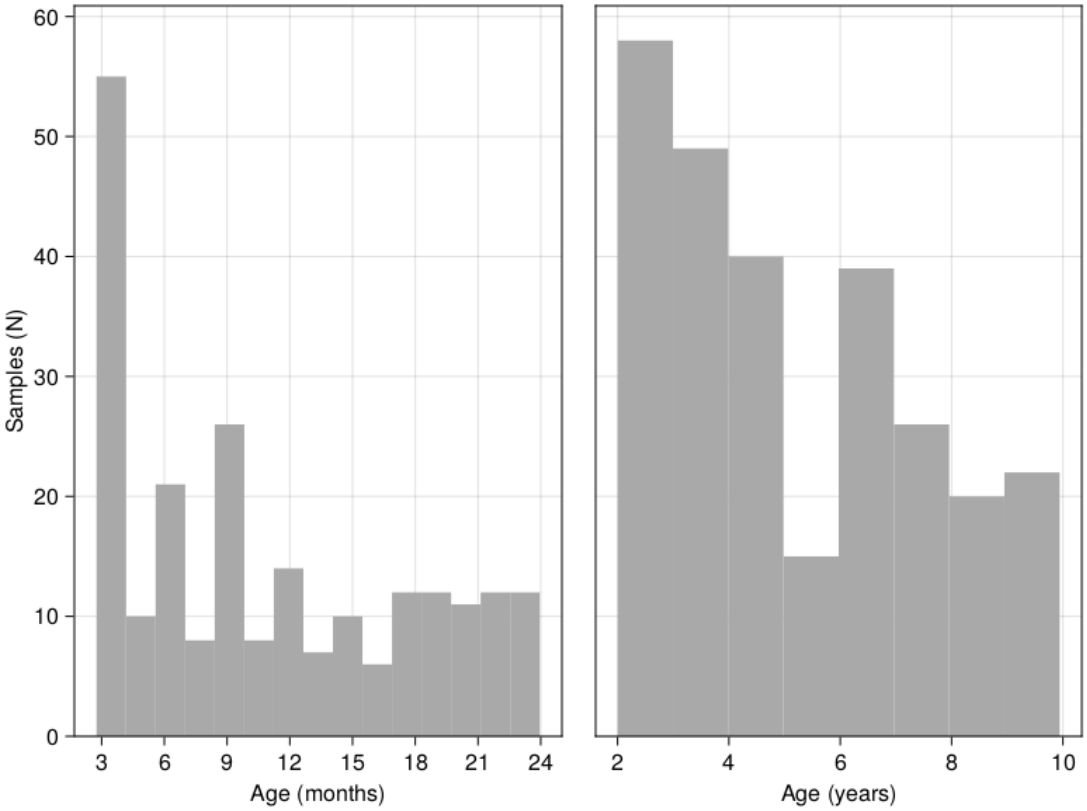


**Figure S1**: Sample collection by age - related to Figure 1A. A histogram showing the number of samples included in this study by the age of the child when the sample was collected.

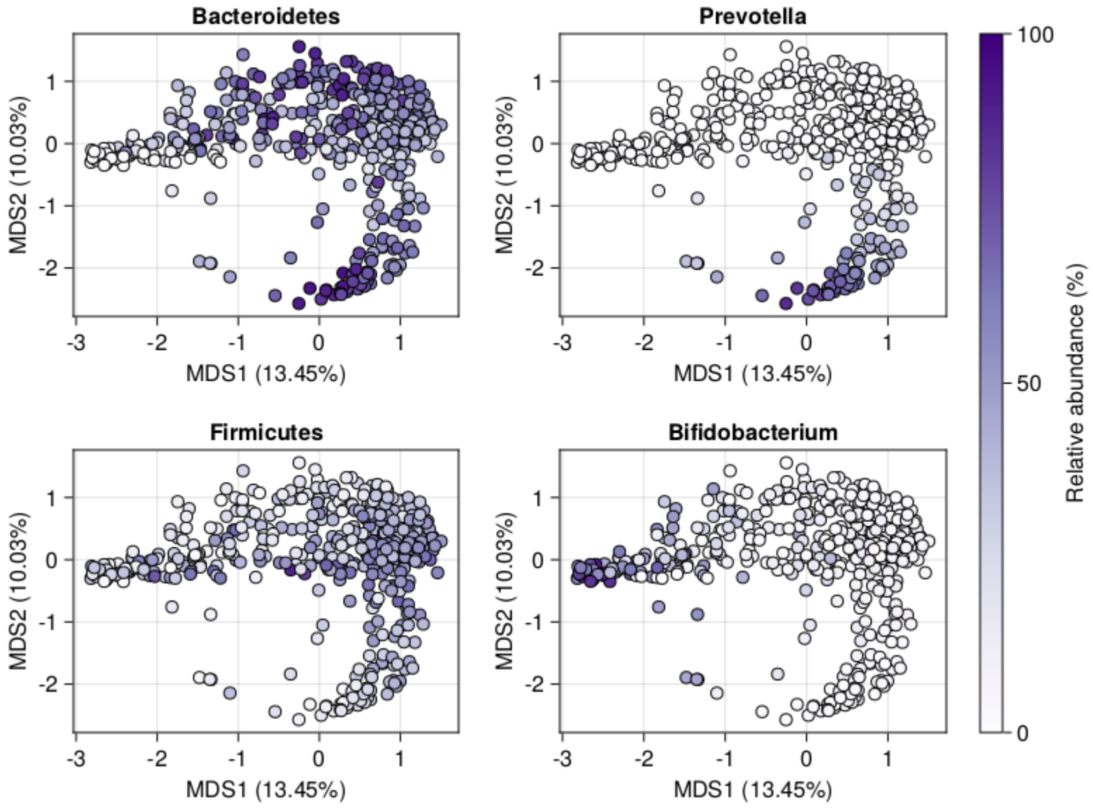


**Figure S2**: Principal coordinates analysis of taxonomic profiles - related to Figure 1C. PCoAs are colored by the relative abundance per-sample of major phyla (Bacteroidetes, top left; Firmicutes, bottom left) and genera (*Prevotella*, top right; *Bifidobacterium*, bottom right).

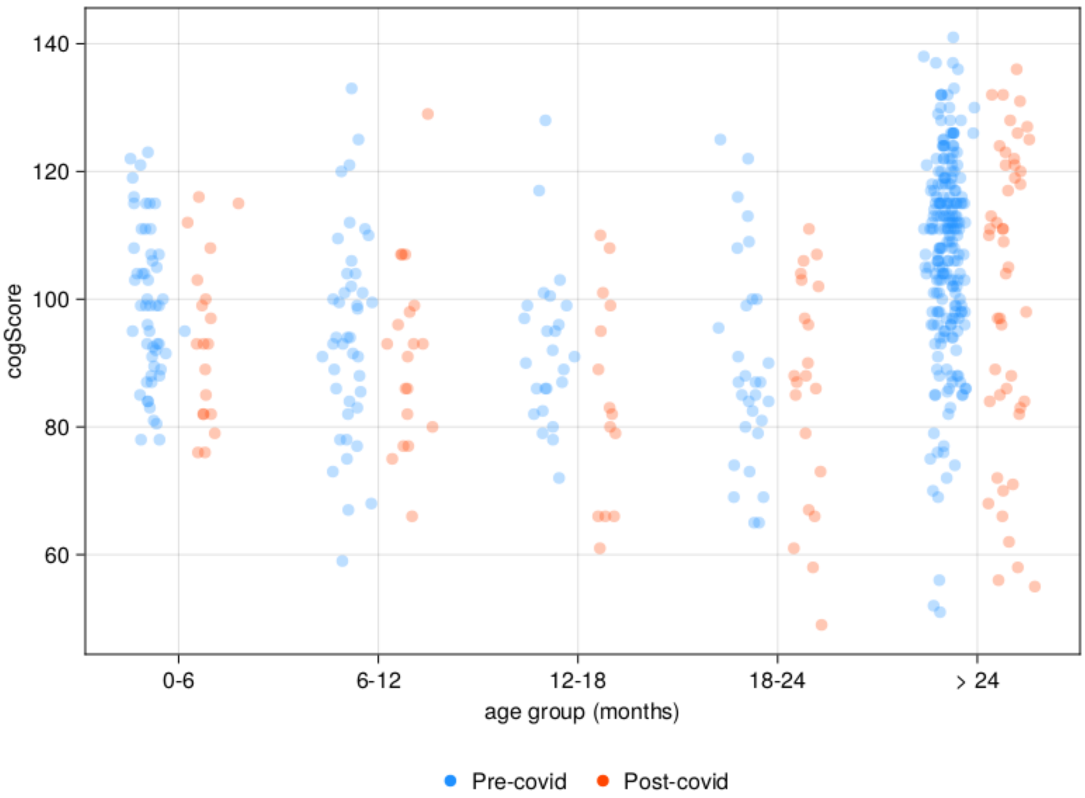


**Figure S3**: Cognitive assessment scores based on age and COVID-19. Scores in red were collected after March of 2020.

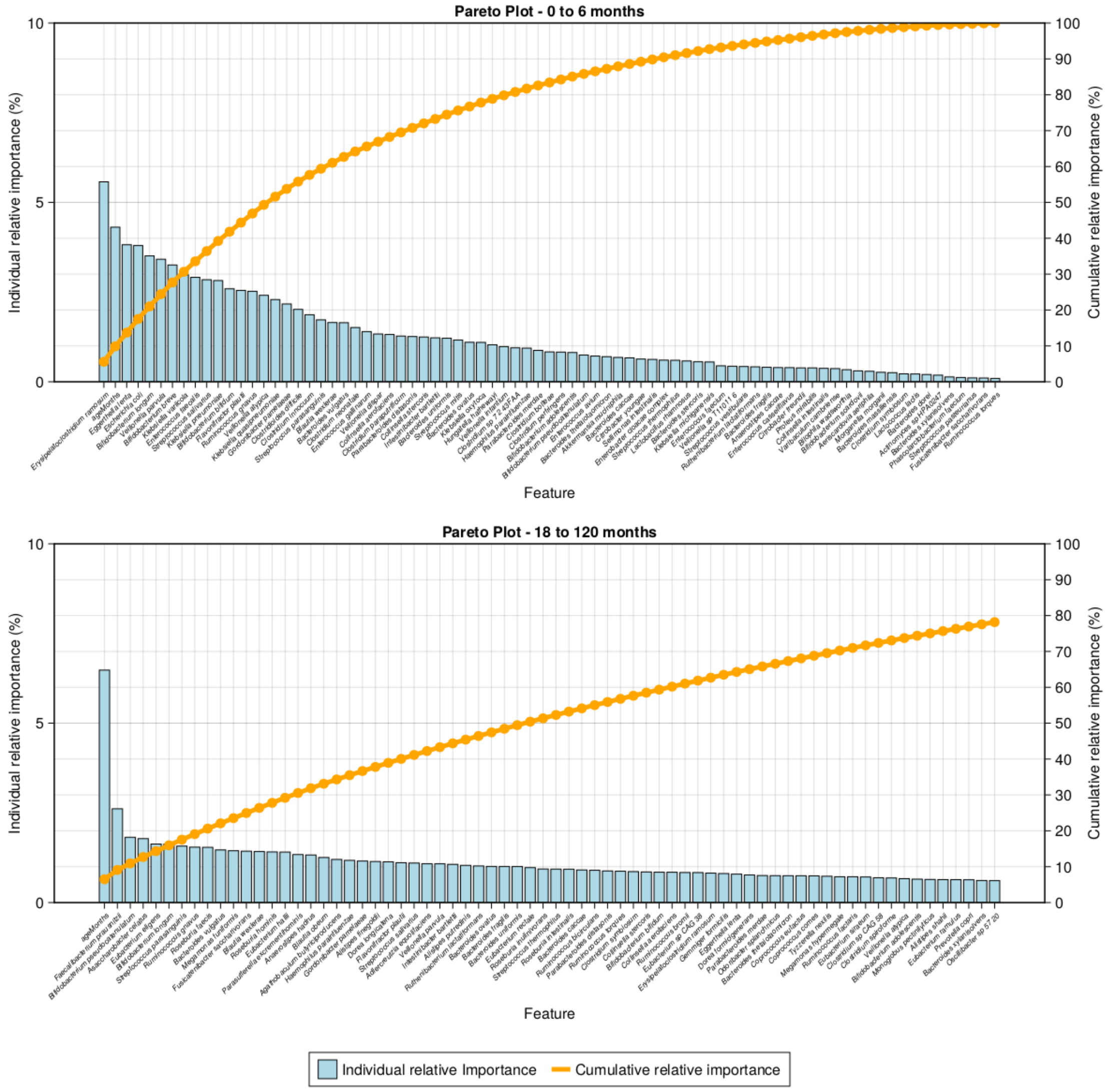


**Figure S4**: Pareto plots for RFs of cognitive function - related to Figure 3C-D. The cumulative importance (orange line), as well as ranked importance (blue bars) of features (species) in models of cogntive performance score in children from birth to 6 months (top) or in children from 18 to 120 months (bottom).

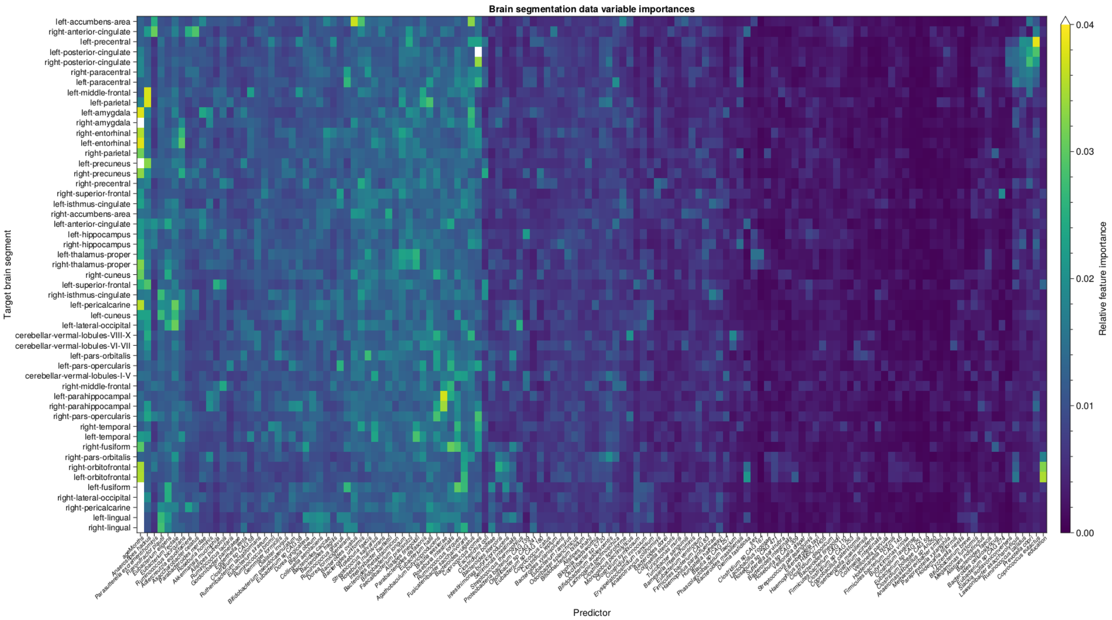


**Figure S5**: Heatmap of feature importances for each brain region - related to Figure 4B. This heatmap contains all features included in models (including age) and all brain segments. Importance values over 0.04 are colored white.

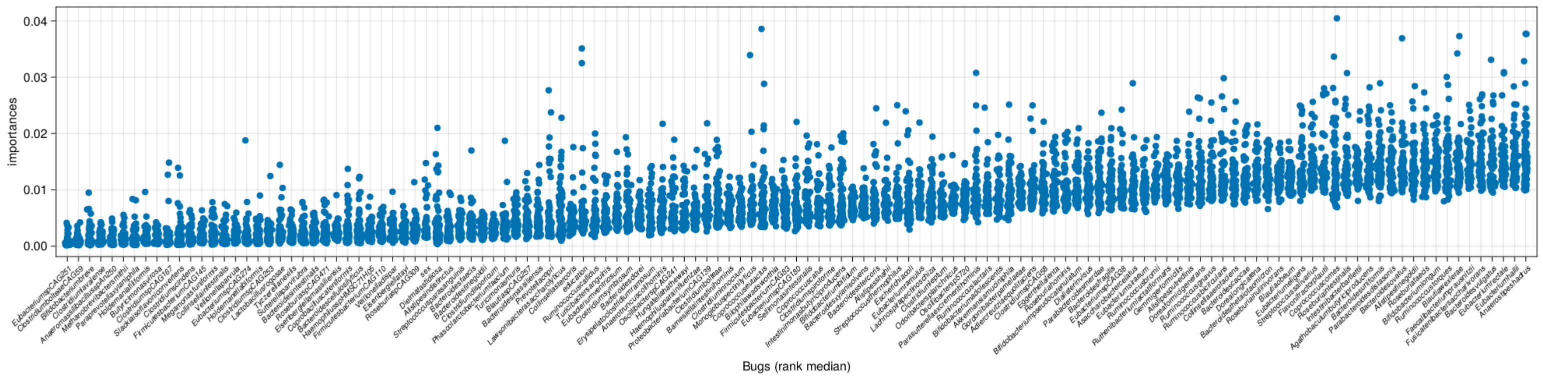


**Figure S6**: Species feature importances in brain region RFs - related to Figure 4C. All species (plus sex and maternal education) feature importances in each brain region model. X-axis is rank-ordered by median importance across all brain region models.

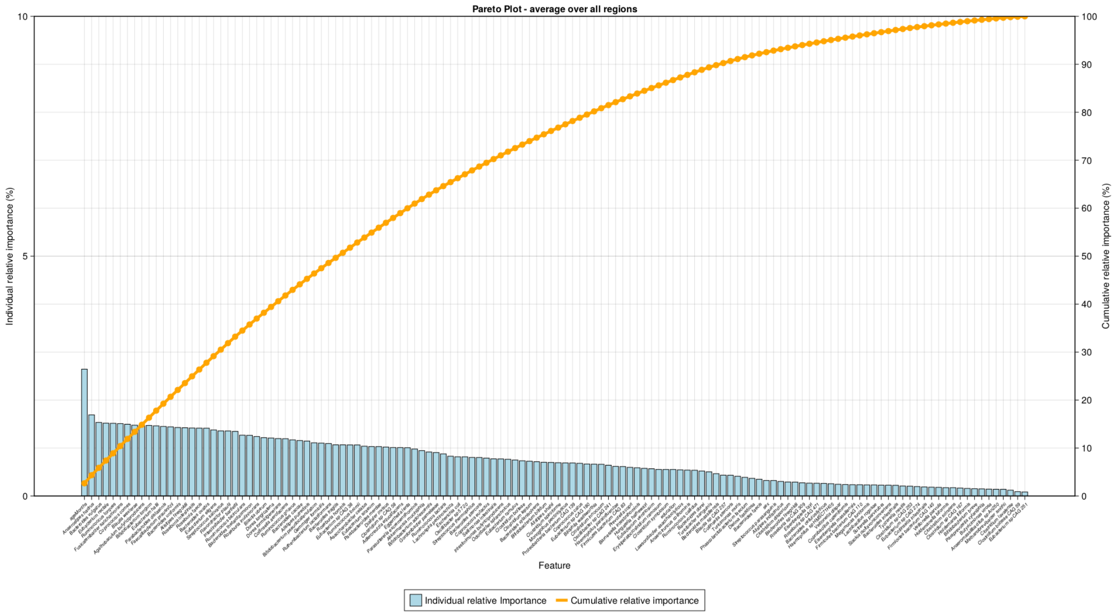


**Figure S7**: Pareto plots for RFs of cognitive function - related to Figure 4. The cumulative average importance (orange line), as well as ranked average importance (blue bars) of features (species) in models of brain region size.

**Table S1A**: Feature importances for RF models of cognitive performance based on taxonomic profiles in children under 6 months old.

| Rank | Variable | Average fitness-weighted importance | Relative fitness-weighted importance | Rank-cumulative relative importance |
|---|---|---|---|---|
| 1 | *Erysipelatoclostridium ramosum* | 0.00122 | 5.57 % | 5.57 % |
| 2 | *ageMonths* | 0.00095 | 4.31 % | 9.88 % |
| 3 | *Eggerthella lenta* | 0.00084 | 3.82 % | 13.7 % |
| 4 | *Escherichia coli* | 0.00083 | 3.8 % | 17.5 % |
| 5 | *Bifidobacterium longum* | 0.00077 | 3.51 % | 21.01 % |
| 6 | *Veillonella parvula* | 0.00075 | 3.42 % | 24.43 % |
| 7 | *Bifidobacterium breve* | 0.00072 | 3.26 % | 27.68 % |
| 8 | *Klebsiella variicola* | 0.00065 | 2.98 % | 30.66 % |
| 9 | *Enterococcus faecalis* | 0.00064 | 2.91 % | 33.57 % |
| 10 | *Streptococcus salivarius* | 0.00062 | 2.84 % | 36.42 % |
| 11 | *Klebsiella pneumoniae* | 0.00062 | 2.82 % | 39.24 % |
| 12 | *Bifidobacterium bifidum* | 0.00057 | 2.59 % | 41.83 % |
| 13 | *Flavonifractor plautii* | 0.00056 | 2.55 % | 44.38 % |
| 14 | *Ruminococcus gnavus* | 0.00055 | 2.52 % | 46.9 % |
| 15 | *Veillonella atypica* | 0.00053 | 2.41 % | 49.31 % |
| 16 | *Klebsiella quasipneumoniae* | 0.0005 | 2.29 % | 51.6 % |
| 17 | *Gordonibacter pamelaeae* | 0.00048 | 2.17 % | 53.77 % |
| 18 | *Clostridioides difficile* | 0.00044 | 2.02 % | 55.79 % |
| 19 | *Clostridium innocuum* | 0.00041 | 1.87 % | 57.66 % |
| 20 | *Streptococcus parasanguinis* | 0.00038 | 1.73 % | 59.38 % |
| 21 | *Blautia wexlerae* | 0.00036 | 1.65 % | 61.03 % |
| 22 | *Bacteroides vulgatus* | 0.00036 | 1.65 % | 62.68 % |
| 23 | *Clostridium neonatale* | 0.00033 | 1.51 % | 64.19 % |
| 24 | *Enterococcus gallinarum* | 0.00031 | 1.4 % | 65.59 % |
| 25 | *Veillonella dispar* | 0.00029 | 1.33 % | 66.92 % |
| 26 | *Collinsella aerofaciens* | 0.00029 | 1.32 % | 68.24 % |
| 27 | *Clostridium paraputrificum* | 0.00028 | 1.27 % | 69.51 % |
| 28 | *Parabacteroides distasonis* | 0.00028 | 1.26 % | 70.78 % |
| 29 | *Collinsella stercoris* | 0.00027 | 1.25 % | 72.02 % |
| 30 | *Intestinibacter bartlettii* | 0.00027 | 1.22 % | 73.25 % |
| 31 | *Bacteroides uniformis* | 0.00027 | 1.21 % | 74.46 % |
| 32 | *Streptococcus mitis* | 0.00026 | 1.16 % | 75.63 % |
| 33 | *Bacteroides ovatus* | 0.00024 | 1.1 % | 76.73 % |
| 34 | *Klebsiella oxytoca* | 0.00024 | 1.1 % | 77.82 % |
| 35 | *Hungatella hathewayi* | 0.00023 | 1.03 % | 78.86 % |
| 36 | *Veillonella infantium* | 0.00022 | 0.98 % | 79.84 % |
| 37 | *Clostridium sp 7 2 43FAA* | 0.00021 | 0.95 % | 80.78 % |
| 38 | *Haemophilus parainfluenzae* | 0.00021 | 0.94 % | 81.72 % |
| 39 | *Parabacteroides merdae* | 0.00019 | 0.88 % | 82.6 % |
| 40 | *Clostridium bolteae* | 0.00018 | 0.83 % | 83.43 % |
| 41 | *Clostridium perfringens* | 0.00018 | 0.83 % | 84.26 % |
| 42 | *Bifidobacterium adolescentis* | 0.00018 | 0.82 % | 85.07 % |
| 43 | *Bifidobacterium pseudocatenulatum* | 0.00016 | 0.75 % | 85.82 % |
| 44 | *Enterococcus avium* | 0.00016 | 0.72 % | 86.53 % |
| 45 | *Bacteroides thetaiotaomicron* | 0.00015 | 0.7 % | 87.23 % |
| 46 | *Akkermansia muciniphila* | 0.00015 | 0.68 % | 87.91 % |
| 47 | *Bacteroides caccae* | 0.00015 | 0.67 % | 88.58 % |
| 48 | *Citrobacter youngae* | 0.00014 | 0.63 % | 89.21 % |
| 49 | *Sellimonas intestinalis* | 0.00014 | 0.62 % | 89.84 % |
| 50 | *Enterobacter cloacae complex* | 0.00013 | 0.6 % | 90.44 % |
| 51 | *Streptococcus thermophilus* | 0.00013 | 0.6 % | 91.04 % |
| 52 | *Lactobacillus rhamnosus* | 0.00013 | 0.58 % | 91.62 % |
| 53 | *Bacteroides stercoris* | 0.00012 | 0.56 % | 92.18 % |
| 54 | *Klebsiella michiganensis* | 0.00012 | 0.55 % | 92.73 % |
| 55 | *Enterococcus faecium* | 0.0001 | 0.45 % | 93.18 % |
| 56 | *Veillonella sp T11011 6* | 0.0001 | 0.43 % | 93.61 % |
| 57 | *Streptococcus vestibularis* | 9.0e-5 | 0.43 % | 94.04 % |
| 58 | *Ruthenibacterium lactatiformans* | 9.0e-5 | 0.42 % | 94.45 % |
| 59 | *Bacteroides fragilis* | 9.0e-5 | 0.41 % | 94.86 % |
| 60 | *Anaerostipes caccae* | 9.0e-5 | 0.4 % | 95.25 % |
| 61 | *Enterococcus casseliflavus* | 9.0e-5 | 0.39 % | 95.65 % |
| 62 | *Citrobacter freundii* | 8.0e-5 | 0.39 % | 96.03 % |
| 63 | *Proteus mirabilis* | 8.0e-5 | 0.39 % | 96.42 % |
| 64 | *Collinsella intestinalis* | 8.0e-5 | 0.37 % | 96.8 % |
| 65 | *Varibaculum cambriense* | 8.0e-5 | 0.37 % | 97.16 % |
| 66 | *Bilophila wadsworthia* | 7.0e-5 | 0.34 % | 97.5 % |
| 67 | *Bifidobacterium scardovii* | 7.0e-5 | 0.3 % | 97.8 % |
| 68 | *Aeriscardovia aeriphila* | 6.0e-5 | 0.29 % | 98.1 % |
| 69 | *Morganella morganii* | 6.0e-5 | 0.26 % | 98.36 % |
| 70 | *Bacteroides massiliensis* | 6.0e-5 | 0.25 % | 98.61 % |
| 71 | *Clostridium symbiosum* | 5.0e-5 | 0.22 % | 98.84 % |
| 72 | *Lactococcus lactis* | 5.0e-5 | 0.22 % | 99.05 % |
| 73 | *Bacteroides dorei* | 4.0e-5 | 0.2 % | 99.26 % |
| 74 | *Actinomyces sp HPA0247* | 4.0e-5 | 0.19 % | 99.44 % |
| 75 | *Bacteroides xylanisolvens* | 3.0e-5 | 0.14 % | 99.58 % |
| 76 | *Phascolarctobacterium faecium* | 3.0e-5 | 0.12 % | 99.7 % |
| 77 | *Streptococcus pasteurianus* | 2.0e-5 | 0.11 % | 99.8 % |
| 78 | *Fusicatenibacter saccharivorans* | 2.0e-5 | 0.1 % | 99.91 % |
| 79 | *Ruminococcus torques* | 2.0e-5 | 0.09 % | 100.0 % |

**Table S1B**: Feature importances for RF models of cognitive performance based on taxonomic profiles in children over 18 months old.

| Rank | Variable | Average fitness-weighted importance | Relative fitness-weighted importance | Rank-cumulative relative importance |
|---|---|---|---|---|
| 1 | *ageMonths* | 0.02129 | 6.48 % | 6.48 % |
| 2 | *Faecalibacterium prausnitzii* | 0.00859 | 2.61 % | 9.09 % |
| 3 | *Bifidobacterium pseudocatenulatum* | 0.00598 | 1.82 % | 10.91 % |
| 4 | *Asaccharobacter celatus* | 0.00586 | 1.78 % | 12.7 % |
| 5 | *Eubacterium eligens* | 0.00536 | 1.63 % | 14.33 % |
| 6 | *Bifidobacterium longum* | 0.00536 | 1.63 % | 15.96 % |
| 7 | *Streptococcus parasanguinis* | 0.00517 | 1.57 % | 17.53 % |
| 8 | *Ruminococcus gnavus* | 0.00507 | 1.54 % | 19.08 % |
| 9 | *Roseburia faecis* | 0.00506 | 1.54 % | 20.62 % |
| 10 | *Bacteroides vulgatus* | 0.00482 | 1.47 % | 22.08 % |
| 11 | *Megamonas funiformis* | 0.00475 | 1.45 % | 23.53 % |
| 12 | *Fusicatenibacter saccharivorans* | 0.00471 | 1.43 % | 24.96 % |
| 13 | *Blautia wexlerae* | 0.00468 | 1.42 % | 26.38 % |
| 14 | *Roseburia hominis* | 0.00464 | 1.41 % | 27.79 % |
| 15 | *Eubacterium hallii* | 0.00462 | 1.41 % | 29.2 % |
| 16 | *Parasutterella excrementihominis* | 0.0044 | 1.34 % | 30.54 % |
| 17 | *Anaerostipes hadrus* | 0.00436 | 1.33 % | 31.87 % |
| 18 | *Blautia obeum* | 0.00413 | 1.26 % | 33.12 % |
| 19 | *Agathobaculum butyriciproducens* | 0.00395 | 1.2 % | 34.32 % |
| 20 | *Haemophilus parainfluenzae* | 0.00386 | 1.18 % | 35.5 % |
| 21 | *Gordonibacter pamelaeae* | 0.0038 | 1.16 % | 36.66 % |
| 22 | *Alistipes finegoldii* | 0.00376 | 1.14 % | 37.8 % |
| 23 | *Dorea longicatena* | 0.00373 | 1.14 % | 38.94 % |
| 24 | *Flavonifractor plautii* | 0.00365 | 1.11 % | 40.05 % |
| 25 | *Streptococcus salivarius* | 0.00362 | 1.1 % | 41.15 % |
| 26 | *Adlercreutzia equolifaciens* | 0.00355 | 1.08 % | 42.23 % |
| 27 | *Veillonella parvula* | 0.00355 | 1.08 % | 43.31 % |
| 28 | *Intestinibacter bartlettii* | 0.0035 | 1.07 % | 44.38 % |
| 29 | *Alistipes putredinis* | 0.00341 | 1.04 % | 45.41 % |
| 30 | *Ruthenibacterium lactatiformans* | 0.00335 | 1.02 % | 46.43 % |
| 31 | *Bacteroides ovatus* | 0.00331 | 1.01 % | 47.44 % |
| 32 | *Bacteroides fragilis* | 0.00331 | 1.01 % | 48.45 % |
| 33 | *Bacteroides uniformis* | 0.0033 | 1.0 % | 49.45 % |
| 34 | *Eubacterium rectale* | 0.0032 | 0.97 % | 50.43 % |
| 35 | *Roseburia inulinivorans* | 0.00307 | 0.94 % | 51.36 % |
| 36 | *Streptococcus thermophilus* | 0.00306 | 0.93 % | 52.29 % |
| 37 | *Roseburia intestinalis* | 0.00305 | 0.93 % | 53.22 % |
| 38 | *Bacteroides caccae* | 0.00298 | 0.91 % | 54.13 % |
| 39 | *Ruminococcus bicirculans* | 0.00296 | 0.9 % | 55.03 % |
| 40 | *Parabacteroides distasonis* | 0.00289 | 0.88 % | 55.91 % |
| 41 | *Ruminococcus torques* | 0.00287 | 0.87 % | 56.78 % |
| 42 | *Clostridium symbiosum* | 0.00283 | 0.86 % | 57.65 % |
| 43 | *Collinsella stercoris* | 0.00281 | 0.85 % | 58.5 % |
| 44 | *Bifidobacterium bifidum* | 0.00278 | 0.85 % | 59.35 % |
| 45 | *Collinsella aerofaciens* | 0.00278 | 0.85 % | 60.19 % |
| 46 | *Ruminococcus bromii* | 0.00275 | 0.84 % | 61.03 % |
| 47 | *Eubacterium sp CAG 38* | 0.00275 | 0.84 % | 61.87 % |
| 48 | *Erysipelatoclostridium ramosum* | 0.0027 | 0.82 % | 62.69 % |
| 49 | *Gemmiger formicilis* | 0.00266 | 0.81 % | 63.5 % |
| 50 | *Eggerthella lenta* | 0.00261 | 0.79 % | 64.29 % |
| 51 | *Dorea formicigenerans* | 0.00253 | 0.77 % | 65.06 % |
| 52 | *Parabacteroides merdae* | 0.00247 | 0.75 % | 65.81 % |
| 53 | *Odoribacter splanchnicus* | 0.00246 | 0.75 % | 66.56 % |
| 54 | *Bacteroides thetaiotaomicron* | 0.00246 | 0.75 % | 67.31 % |
| 55 | *Coprococcus eutactus* | 0.00246 | 0.75 % | 68.06 % |
| 56 | *Coprococcus comes* | 0.00245 | 0.74 % | 68.8 % |
| 57 | *Tyzzerella nexilis* | 0.00241 | 0.73 % | 69.54 % |
| 58 | *Megamonas hypermegale* | 0.00236 | 0.72 % | 70.25 % |
| 59 | *Ruminococcus lactaris* | 0.00236 | 0.72 % | 70.97 % |
| 60 | *Eubacterium siraeum* | 0.00234 | 0.71 % | 71.69 % |
| 61 | *Clostridium sp CAG 58* | 0.00227 | 0.69 % | 72.38 % |
| 62 | *Clostridium spiroforme* | 0.00225 | 0.69 % | 73.06 % |
| 63 | *Veillonella atypica* | 0.00219 | 0.67 % | 73.73 % |
| 64 | *Bifidobacterium adolescentis* | 0.00214 | 0.65 % | 74.38 % |
| 65 | *Monoglobus pectinilyticus* | 0.00212 | 0.64 % | 75.03 % |
| 66 | *Alistipes shahii* | 0.0021 | 0.64 % | 75.67 % |
| 67 | *Eubacterium ramulus* | 0.0021 | 0.64 % | 76.31 % |
| 68 | *Prevotella copri* | 0.00209 | 0.64 % | 76.94 % |
| 69 | *Bacteroides xylanisolvens* | 0.00202 | 0.62 % | 77.56 % |
| 70 | *Oscillibacter sp 57 20* | 0.00201 | 0.61 % | 78.17 % |
| 71 | *Blautia sp CAG 257* | 0.00198 | 0.6 % | 78.77 % |
| 72 | *Sutterella parvirubra* | 0.00198 | 0.6 % | 79.38 % |
| 73 | *Akkermansia muciniphila* | 0.00192 | 0.58 % | 79.96 % |
| 74 | *Clostridium bolteae* | 0.00186 | 0.57 % | 80.53 % |
| 75 | *Clostridium leptum* | 0.00185 | 0.56 % | 81.09 % |
| 76 | *Lachnospira pectinoschiza* | 0.00185 | 0.56 % | 81.66 % |
| 77 | *Bacteroides stercoris* | 0.00185 | 0.56 % | 82.22 % |
| 78 | *Clostridium innocuum* | 0.00184 | 0.56 % | 82.78 % |
| 79 | *Escherichia coli* | 0.00183 | 0.56 % | 83.33 % |
| 80 | *Eubacterium sp CAG 180* | 0.00179 | 0.55 % | 83.88 % |
| 81 | *Intestinimonas butyriciproducens* | 0.00179 | 0.55 % | 84.42 % |
| 82 | *Dialister invisus* | 0.00177 | 0.54 % | 84.96 % |
| 83 | *Anaerotruncus colihominis* | 0.00171 | 0.52 % | 85.48 % |
| 84 | *Bifidobacterium breve* | 0.00168 | 0.51 % | 85.99 % |
| 85 | *Sellimonas intestinalis* | 0.00166 | 0.51 % | 86.5 % |
| 86 | *Veillonella infantium* | 0.00162 | 0.49 % | 86.99 % |
| 87 | *Bacteroides dorei* | 0.00161 | 0.49 % | 87.48 % |
| 88 | *Proteobacteria bacterium CAG 139* | 0.0016 | 0.49 % | 87.97 % |
| 89 | *Megamonas funiformis CAG 377* | 0.00152 | 0.46 % | 88.43 % |
| 90 | *Bilophila wadsworthia* | 0.00152 | 0.46 % | 88.89 % |
| 91 | *Hungatella hathewayi* | 0.00145 | 0.44 % | 89.33 % |
| 92 | *Veillonella dispar* | 0.00144 | 0.44 % | 89.77 % |
| 93 | *Turicimonas muris* | 0.00141 | 0.43 % | 90.2 % |
| 94 | *Roseburia sp CAG 471* | 0.00141 | 0.43 % | 90.63 % |
| 95 | *Eisenbergiella massiliensis* | 0.00137 | 0.42 % | 91.05 % |
| 96 | *Firmicutes bacterium CAG 83* | 0.00127 | 0.39 % | 91.44 % |
| 97 | *Barnesiella intestinihominis* | 0.00126 | 0.38 % | 91.82 % |
| 98 | *Turicibacter sanguinis* | 0.00122 | 0.37 % | 92.19 % |
| 99 | *Bifidobacterium catenulatum* | 0.00117 | 0.36 % | 92.55 % |
| 100 | *Bacteroides finegoldii* | 0.00115 | 0.35 % | 92.9 % |
| 100+ | See Table file | ... | ... | ... |

**Table S2**: RF model performances predicting cortical and subcortical brain regions.

| Segment | Mean absolute proportional error (MAPE) | Correlation coefficient (R) |
|---|---|---|
| Left temporal | 0.0405 | 0.1278 |
| Right temporal | 0.0393 | 0.0683 |
| Left orbitofrontal | 0.0554 | 0.1862 |
| Right orbitofrontal | 0.0614 | 0.1325 |
| Left parietal | 0.0352 | 0.1767 |
| Right parietal | 0.0451 | -0.1471 |
| Left middle frontal | 0.0521 | -0.0002 |
| Right middle frontal | 0.0385 | 0.0829 |
| Left anterior cingulate | 0.0685 | 0.0917 |
| Right anterior cingulate | 0.0739 | 0.2447 |
| Left lateral occipital | 0.0622 | 0.2237 |
| Right lateral occipital | 0.0544 | 0.2208 |
| Left cerebellum white matter | 0.0667 | 0.4132 |
| Right cerebellum white matter | 0.0698 | 0.4344 |
| Left thalamus proper | 0.0534 | 0.1696 |
| Right thalamus proper | 0.0499 | 0.1616 |
| Left caudate | 0.099 | 0.0287 |
| Right caudate | 0.083 | -0.01 |
| Left putamen | 0.0592 | 0.1541 |
| Right putamen | 0.0658 | 0.0107 |
| Left pallidum | 0.0641 | 0.2967 |
| Right pallidum | 0.0613 | 0.2644 |
| Left hippocampus | 0.0541 | 0.0749 |
| Right hippocampus | 0.0593 | -0.0937 |
| Left amygdala | 0.0632 | 0.2879 |
| Right amygdala | 0.0704 | 0.1787 |
| Left accumbens area | 0.0941 | 0.2877 |
| Right accumbens area | 0.1024 | -0.0406 |
| Left basal forebrain | 0.2059 | 0.1544 |
| Right basal forebrain | 0.1169 | 0.1234 |
| Left cuneus | 0.0816 | 0.2345 |
| Right cuneus | 0.075 | 0.0647 |
| Left entorhinal | 0.0798 | 0.1904 |
| Right entorhinal | 0.0822 | 0.0993 |
| Left fusiform | 0.0572 | 0.3331 |
| Right fusiform | 0.053 | 0.2482 |
| Left isthmus cingulate | 0.0774 | -0.0409 |
| Right isthmus cingulate | 0.0854 | 0.0735 |
| Left lingual | 0.0688 | 0.4212 |
| Right lingual | 0.0719 | 0.4336 |
| Left parahippocampal | 0.0576 | 0.1535 |
| Right parahippocampal | 0.0602 | 0.1341 |
| Left paracentral | 0.0781 | 0.1309 |
| Right paracentral | 0.084 | 0.0231 |
| Left pars opercularis | 0.057 | 0.1464 |
| Right pars opercularis | 0.0595 | 0.0301 |
| Left pars orbitalis | 0.0821 | 0.0032 |
| Right pars orbitalis | 0.1118 | 0.0235 |
| Left pars triangularis | 0.0633 | 0.1743 |
| Right pars triangularis | 0.0762 | 0.0434 |
| Left pericalcarine | 0.1105 | 0.1997 |
| Right pericalcarine | 0.0885 | 0.2729 |
| Left postcentral | 0.0516 | 0.1681 |
| Right postcentral | 0.0595 | 0.1349 |
| Left posterior cingulate | 0.1338 | 0.1365 |
| Right posterior cingulate | 0.1394 | 0.0628 |
| Left precentral | 0.0654 | 0.2478 |
| Right precentral | 0.0463 | 0.0656 |
| Left precuneus | 0.0561 | 0.1443 |
| Right precuneus | 0.0685 | 0.1499 |
| Left superior frontal | 0.0398 | 0.1008 |
| Right superior frontal | 0.0436 | -0.0961 |
| Left supramarginal | 0.0668 | 0.0611 |
| Right supramarginal | 0.0682 | -0.0008 |
| Left insula | 0.0571 | -0.119 |
| Right insula | 0.0562 | 0.0771 |
| Cerebellar vermal lobules I V | 0.0846 | 0.1562 |
| Cerebellar vermal lobules VI VII | 0.0813 | 0.006 |
| Cerebellar vermal lobules VIII X | 0.0825 | 0.0781 |
| Brain stem | 0.0766 | 0.6047 |
| CSF | 0.1624 | 0.0713 |

**Table S3**: Experimental design and input composition for Random Forest experiments.

| Input set | Age bracket | Microbiome encoding type | Demographics Provided? (sex, education) |
|---|---|---|---|
| 1 | 0 to 6 months | Not provided | yes |
| 2 | | Taxonomic profile | no |
| 3 | | | yes |
| 4 | | Functional Profile (ECs) | no |
| 5 | | | yes |
| 6 | 18 to 120 months | Not provided | yes |
| 7 | | Taxonomic profile | no |
| 8 | | | yes |
| 9 | | Functional Profile (ECs) | no |
| 10 | | | yes |
